# Does embedding impact classification performance ? A small sample perspective

**DOI:** 10.64898/2026.11.24.768658

**Authors:** Marek Sztuka, Joanna Szyda

**Author notes:** Correspondence to: Joanna Szyda, Department of Biomedical Engineering, Wroclaw University of Science and Technology, Plac Grunwaldzki 13, 50-377 Wroclaw, Poland.

## Abstract

Deep learning (DL) methods can address many challenges posed by high-dimensional and sparse microbiome data. In this study we applied and compared deep (DNN) and shallow (SNN) feedforward neural networks combined with five dimensionality-reduction techniques: Principal Component Analysis (PCA), Singular Value Decomposition (SVD), Non-negative Matrix Factorization (NMF), Autoencoder based Embeddings (AE), and convolutional layers (Conv). To establish baseline classification performance, the XGBoost was used, as it represents a robust machine learning based classifier with a relatively low number of hyperparameters. The task was to classify effective microorganism (EM) treatments based on intestine bacterial family abundance in common carp. In particular, the experiment involved 25 ponds that were divided into experimental groups. In each pond, the intestinal microbiome of five individuals was determined by sequencing two hypervariable regions of the gene encoding the 16s rRNA gene sequencing. The sequence reads were processed by QIIME2 and taxonomically annotated to the SILVA database, what resulted in 126 annotated families that were used as features in all downstream analyses. Abundance tables from 125 samples, transformed using the Centered Log-Ratio (CLR) method, were used to evaluate ten dimensionality-reduction – Deep learning configurations considering five-, three-, and two-classification scenarios. The DL models were optimized using cross-validation. To combat the overfitting that was present due to sparsity and small sample size, dropout and kernel regularization were implemented. During testing autoencoder embeddings achieved the highest classification accuracy (0.84), exceeding the XGBoost baseline (0.64), while embedding based on convolutional layers significantly reduced performance (0.44). PCA and SVD transformations showed comparable results (0.65), whereas NMF often failed to converge, likely due to incompatibility with CLR-transformed data. Best classification results were obtained for models that implemented AE embedding, it is most likely caused by their ability to capture non-linear patterns in the data. All models showed stable training behavior, with loss functions consistently decreasing across epochs. The SNN models were markedly less computationally intensive, than DNN. AE and NMF based embeddings were the most resource hungry (Wall Clock Time and RAM). Our results show (1): SNN architectures performed slightly worse than DNN, due to the fact of lesser informative capacity of the models. (2): Dimensionality reduction impacts model performance, though not always positively. (3) EM supplementation causes significant changes in microbiome that can be captured by DL architectures. Future work will include feature-importance analysis, integration of environmental and host metadata, exploration of larger datasets and more advanced architectures. As well as validation of model quality on simulated datasets. Overall, these results indicate that DNN architectures combined with AE embeddings have potential to be used as high accuracy classification tools. Those tools when combined with feature importance can help researchers to understand changes made by EM supplementation in a microbial community.

## I. Introduction

Pattern recognition with Deep learning (DL) has a long history of successful applications, tracing back over three decades to the seminal study by LeCun et.al. [1] on handwritten digit classification. Despite these advances, robust pattern extraction for experimental data often remains challenging due to high levels of noise, data sparsity, missing and erroneous measuremests, or even ambiguous outcome labelling. Biological datasets in particular, introduce further complexities, including varying degree of feature correlations, patterns driven only by a small subset of features (or even a single feature. Due to the biological and ethical constraints of data acquisition, biological datasets often exhibit high dimensionality, even though the number of subjects is small. Thus is driven by the discrepancy between a small number of subjects (N) characterised by a large number of features (P). Modern Genome-Wide Association Studies (GWAS) well exemplify this challenge, where phenotypes scored for relatively few individuals are tested for association with genetic variants across the whole genome. For instance, the Million Veteran Program - one of the largest available human cohorts - consists of 635,000 participants (N) described by over 44.3 million genetic polymorphisms (P) [2], illustrating a severe P>>N problem.

In our analysis, we explore approaches to mitigate the P>>N problem by various input embedding strategies, using fish gut microbiome abundance dataset as a case study. This data was selected because, despite growing interest in microbiome analysis, it remains statistically challenging. While the overall data set dimensions are small, its high-dimensionality is due to the discrepancy between the number of sequenced microbial communities (N) and the number of features, here represented by bacterial families (P). Such data poses an evaluation problem not directly because of elevated P>>N, but rather because the fact that the low number of subjects does not provide much information to handle the comparatively large number of features and zero inflated counts in the context of classification. Eloe-Fadrosh et al. (2015) and Ni et al. (2025)[3], [4] evaluate cohorts of only 12 and 20 subjects, respectively, against hundreds of bacterial taxa, and underscoring downsides of state of the art methodology in terms of identification power across feature space with limited sample size. Traditional statistical tests for differential abundance typically compare each taxon independently, which not only ignores taxa co-occurrence dynamics but also inflates type I error of hypotheses testing [5]. To address these limitations, researchers increasingly rely on reduced dimensionality expressed by embedded feature representations derived from either more natural - linear or non-linear transformations. These include deterministic approaches such as Principal Component Analysis (PCA)[6], Singular Value Decomposition (SVD) [7], and Nonnegative Matrix Factorization (NMF)[8] or even stochastic dimensionality reduction methods such as Uniform Manifold Approximation and Projection (UMAP)[9], or t-Distributed Stochastic Neighbor Embedding (t-SNE)[10], for capturing latent patterns in sparse abundance representations [11], [12]. Our study extends those traditional approaches by using DL-based embeddings, expressed by a latent space of Autoencoder architectures [13] or by a 1-dimensional Convolutional Neural Network [14]. All the embedding approaches project high-dimensional microbiome abundance data into a lower-dimensional latent manifold, improving computational efficiency, reducing noise and circumvent algebraic problem of zero-counts. However, they often compromise on biological interpretability, making it difficult to trace which bacterial taxa drive classification outcomes. Such data structures, here represented by the microbiome, are also often used in experimental [ref] or medical context [ref] where a large number of subjects is not available due to ethical or financial reasons.

First conceptualized by McCulloch and Pitts in 1943 [15] and developed over decades, neural networks are computational models that provide an alternative to rigorous statistical modelling. Recently, they have gained in popularity due to the widespread availability of computational resources operating in the parallelized mode and user-friendly implementation frameworks like TensorFlow [16] further facilitated through Keras [17] interface or PyTorch [18]. Unlike traditional tree-based methods (e.g. Random Forests or XGBoost), Deep learning (DL) architectures offer the distinct advantage of end-to-end representation learning. An added value of DL architectures over is their ability to integrate embedding layers directly into a training and to apply regularization techniques such as dropout. These features are particularly effective for handling sparse and noisy data, such as microbiome abundance counts. Furthermore, thanks to processing all features simultaneously, through multiple representation layers, DL models architectures can provide better fit to learn complex, nonlinear relationships between co-occurring bacterial taxa, potentially uncovering biologically meaningful patterns inaccessible than via conventional statistics to conventional statistical and shallow machine learning methods, and thus have a better potential to uncover biologically meaningful patterns.

In this study, we aim to evaluate which dimensionality reduction techniques (DRTs) can best preserve relevant biological information from microbiome data and improve classification performance. The underlying motivation was driven by the fact that, that due to the low number of samples and a zero inflated structure of the data, it may be beneficial to reduce the number of input features without compromising information content. Such a reduced number of features shall allow for better classification performance, as it better represents the variation of the original data and mitigates the curse of dimensionality problem. While we apply this approach to gut microbiome abundance data, the methodology is highly adaptable to other sparse tabular datasets with similar structural properties defined by the small number of subjects and a comparatively large number of features, with the focus on classification with underlaying class ambiguity.

To summarise, this study explores DL-based pattern recognition approaches for pattern recognition and classification strategies in mildly high-dimensional, compositional datasets. We benchmark shallow and deep neural network architectures against random class assignment and against robust traditional baselines implementing the XGBoost approach across various class definitions (five, three, and two classes). Additionally, we examine various embedding techniques to capture data patterns and assess their impact on classification performance. Finally, to assess the robustness and generalizability of our real data results, we validate our findings through a simulation study [19].

## II. The data

The data set used in the study originates from the field experiment investigating the effects of water and feed supplementation with commercial Effective Microorganism (EM) additives on the microbial composition of water, sediment, and fish gut [20]. The common carp (*Cyprinus carpio*) served as the model organism. The experimental design comprised five groups varying in regard to EM supplementation schemes: consisting of water supplements (W1, W2) and feed supplements (F1, F2). Specifically, the design included: a control group (no supplementation), Group 1 (W1 only), Group 2 (W1 and F1), Group 3 (W2 only), and Group 4 (W2 and F2) (Table I). Each group was represented by five ponds. At the end of the experiment, intestinal samples were collected from five fish per pond, yielding 124 microbiome samples (one sample was excluded due to a technical reasons). Taxonomic annotation of microbiome samples was based on sequences of two hypervariable regions of the 16S rRNA gene. Raw reads were pre-processed and taxonomically annotated using the standard QIIME2 [21] pipeline. This resulted in the dataset containing 126 bacterial families (features) across 124 samples - a semi-square data frame that does not exactly express the severe P>>N structure discussed above, but still represents a zero-inflated, count data set with the number of features exceeding the number of subjects. In the next step, the read counts representing each bacterial family were rescaled using Centered Log-Ratio (CLR) [22] transformation, which is a standard choice for abundance data, which in nature has compositional structure [5] as it allows to account for such factors as variable sequencing depth across samples [23]. Note, that in this experiment an additional complexity is added by the fact that the true number of classes remains uncertain, as the efficacy of the supplementation schemes in altering the microbiome composition is unknown.

**TABLE I.** POND SUPPLEMENTATION SCHEME.

|  | **Control<br>5 ponds** | **Group 1<br>5 ponds** | **Group 2<br>5 ponds** | **Group 3<br>5 ponds** | **Group 4<br>5 ponds** |
|---|---|---|---|---|---|
| **Fish feed** | none | none | F1 | none | F2 |
| **Water** | none | W1 | W1 | W2 | W2 |

## III. Methods

### A. Class Definition and Classification Baselines

The first part of this study focuses on evaluating various Deep learning architectures and dimensionality reduction techniques to establish the best approach to classify ponds based on microbiome abundance patterns (Figure 1). In the first step, data was divided into a test set consisting of 25 intestinal samples and a train-validation set of 99 intestinal samples.

**Fig. 1.**
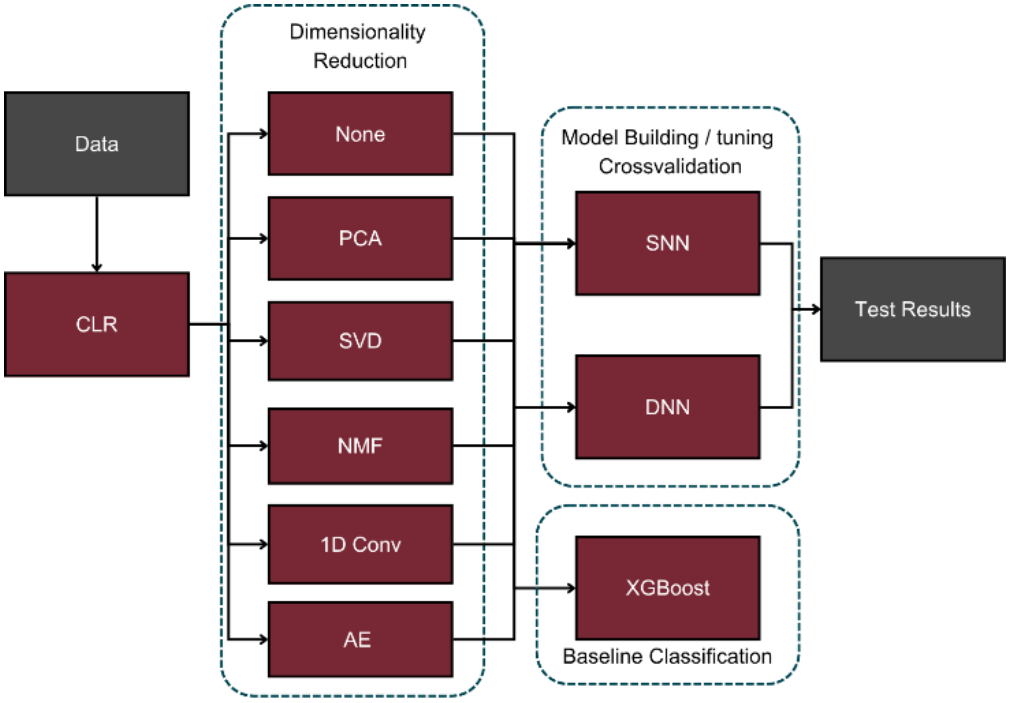
Case study analysis pipeline schema

Given that the initial 5-class split may not reflect a true biological pattern, we additionally evaluated alternative class splits. Specifically, (i) considering the sole effect of feed supplementation vs. the combined effect of water and feed supplementation, the following three classes were considered: the **control group** (no supplementation), the **water supplementation group** (group 1 and group 3), and the **combined supplementation group** (group 2 and group 4) as well as (ii) considering only the effect of feed supplementation; the following two classes were considered: the **group without feed supplementation** (control group, group 1, and group 3) and the **group with feed supplementation** (group 2 and group 4) (Table II). Under a random baseline representing the null hypothesis of no effect of supplementation, expressed by random assignment of samples to classes, the accuracies are calculated as the reciprocal of the number of classes as 0.20 for the 5-class split, 0.33 for the 3-class split, and 0.50 for the 2-class split. Under the alternative hypothesis assuming the effect of supplementation, a machine learning baseline was represented by the XGBoost gradient-boosted decision tree classifier [24], [25]. The model was configured based on trial and error manner with a maximum tree depth of 3 and a learning rate of 0.1. It was trained for 100 iterations.

**TABLE II.** CLASS SPLITS.

|  |  |  |  |  |  |
|---|---|---|---|---|---|
| 5-class-split | Control | Group 1 | Group 2 | Group 3 | Group 4 |
| 3-class-split | Control | Groups 1 and 3 |  | Groups 2 and 4 |  |
| 2-class-split | Groups Control, 1 and 3 |  |  | Groups 2 and 4 |  |

### B. Data embedding

In this study, we evaluated five dimensionality reduction approaches. The first technique was Principal Component Analysis (PCA) [26]. To ensure consistency in the modeling pipeline, the PCA transformation was fitted on the training dataset and subsequently applied to transform both the training and test datasets. The optimal latent dimensionality of PCA was determined by targeting a cumulative explained variance threshold of 90% and subsequently evaluating the classification performance for adjacent cardinalities. The second representation was based on Singular Value Decomposition (SVD) [27]. Like for PCA, SVD was computed exclusively on the training data set and the final number of components was determined based on accuracies estimated for the selection of components adjacent the set defining the 90% of variance. The third strategy was Non-negative Matrix Factorization (NMF) [28] which requires non-negative input space. Consequently, the data set was preprocessed using the *log*(*x* + *e*) transformation, where *e* represents 1*e*^−5^ to eliminate zeroes. 99 latent components were estimated over maximum of 10,000 iterations and subsequently used as the reduced representation of the features.

The fourth approach employed a Deep learning-based embedding using a Autoencoder (AE) [29]. We utilized a symmetrical AE architecture with two fully connected encoder/decoder layers fitting 64 and 32 nodes, employing linear and ReLU activation functions respectively. Latent space in between with 19 nodes corresponding to the 19-dimensional embedding vector used in PCA (Fig. 1). Note that although using 19 nodes may be suboptimal from the perspective of computational efficiency, it was the optimal length of the latent space resulting from the hyperparameter search and was retained because the computational time was not a critical aspect for this small data set. The Mean Squared Error (MSE) loss function was optimized using the Adam optimizer [30] until the loss plateaued which was monitored by Keras early stopping callback, with patience parmeter set to 10. This final network architecture was determined by a manual search of the hyperparameter space of layer depths and the numbers of nodes, using the lowest mean feature reconstruction error as the model selection criterion.

The fifth embedding strategy utilized two one-dimensional Convolutional Neural Networks (1DConv). The first layer comprised 2 filters with a kernel size of 25, followed by a second 1DConv utilizing a single filter with a kernel size of 3. Subsequently, to aggregate convolution output, the resulting feature representations were downsampled using a 1D max-pooling operation with a pool size of 4. Finally, a Dense layer with 64 units and a Flatten layer, which reshaped the output into a one-dimensional vector suited for downstream classification models (Fig. 2). Choice of architecture was dictated by manual hyperparameter tuning with best classification accuracy in mind.

**Fig. 2.**
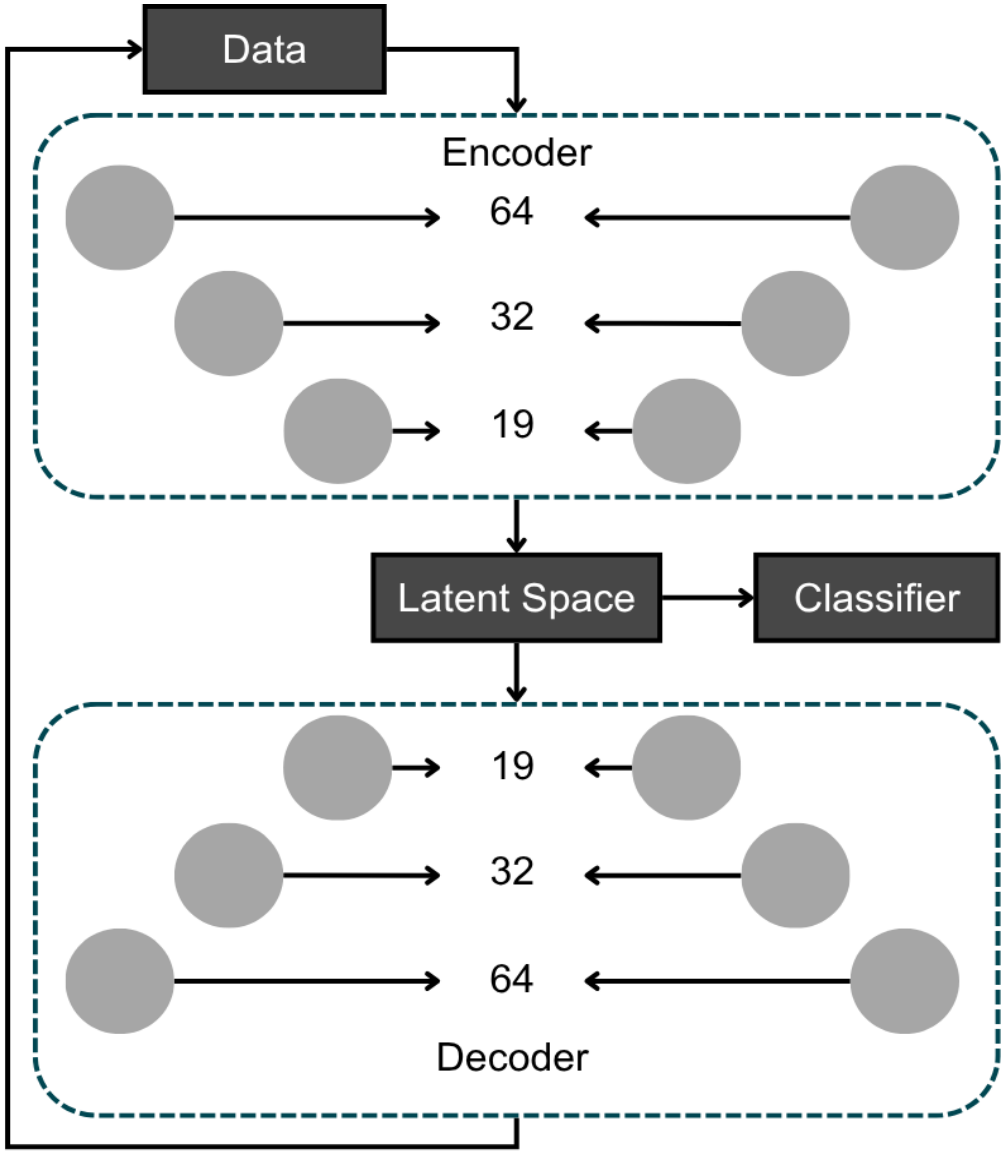
Autoencoder architecture for dimensionality reduction.

**Fig. 3.**
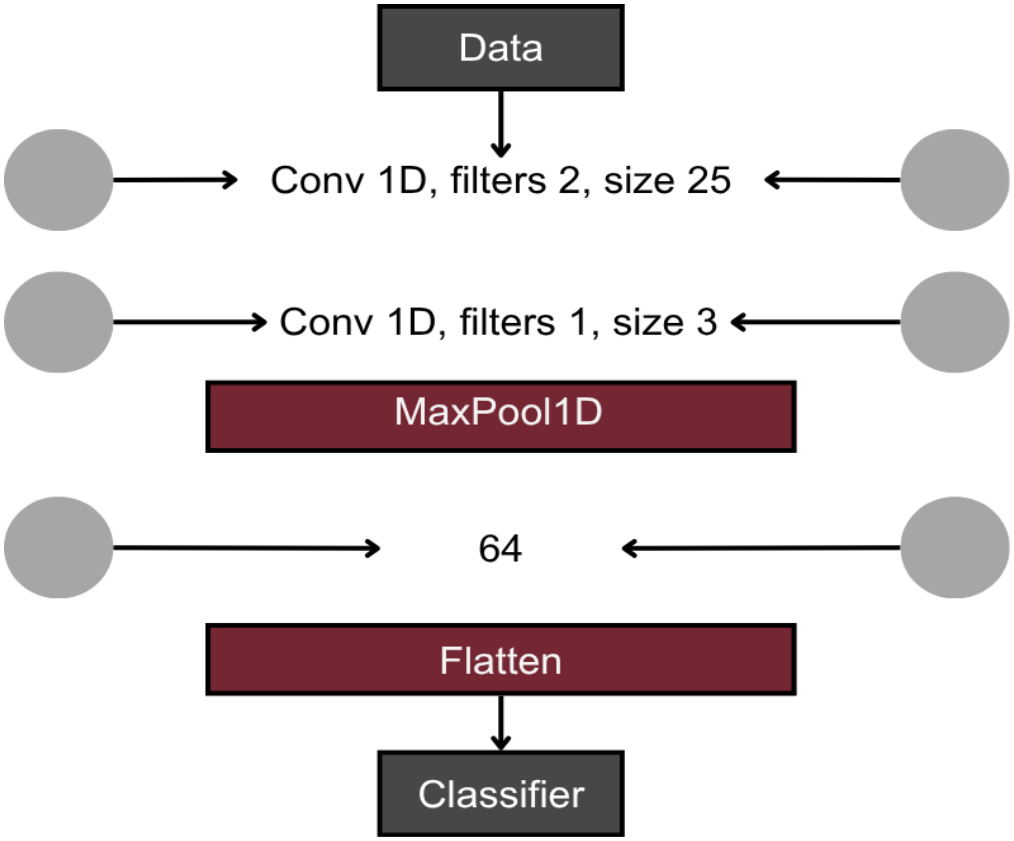
Convolutional dimensionality reduction architecture

### C. Classifiers

The next step of the analysis was to evaluate two feed-forward neural network topologies for classification: a Deep Neural Network (DNN) and a Shallow Neural Network (SNN). Specifically, the DNN architecture comprised of four dense layers with progressively decreasing dimensions (Table III), while SNN represented a minimal configuration aimed to evaluate the simplest possible architecture consisting of a single dense layer with a number of nodes close to the input dimensionality (Table IV). Both architectures shared a standardized training protocol and hyperparameters: the Rectified Linear Unit (ReLU) was used for activation, the Softmax activation was applied in the output layer for multi-class classification, loss was defined as categorical cross-entropy function and was optimized by the Adam algorithm [30]. To mitigate overfitting in this small N situation, both architectures incorporated *L*2 weight regularization on all dense layers, as well as dropout layers with a 0.25 dropout rate, positioned after the first dense layer. The network terminated with an output layer matching the target number of classes (5, 3, or 2).

**TABLE III.**
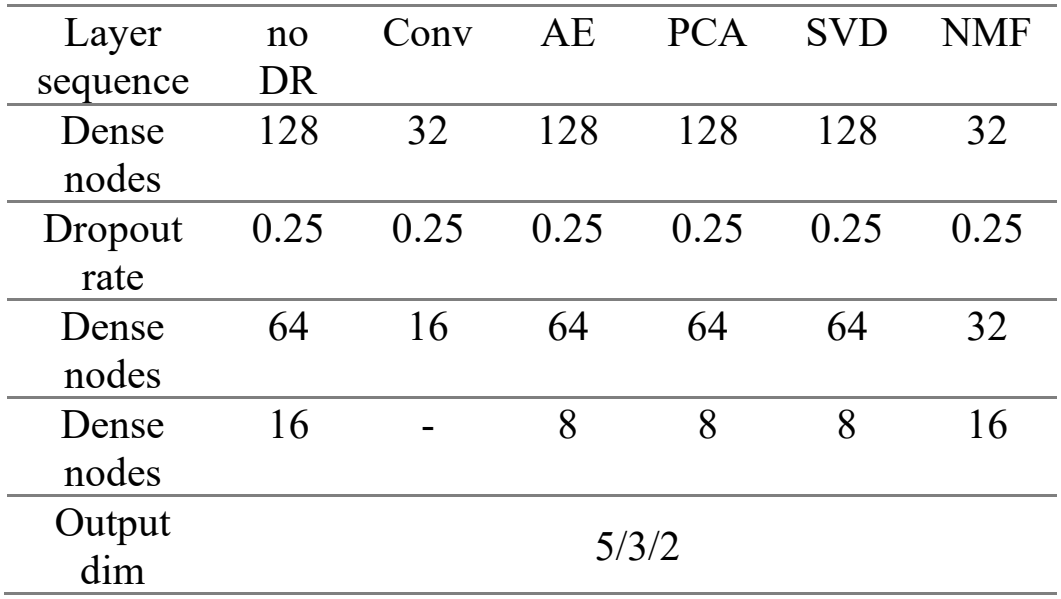
DNN ARCHITECTURE.

**TABLE IV.**
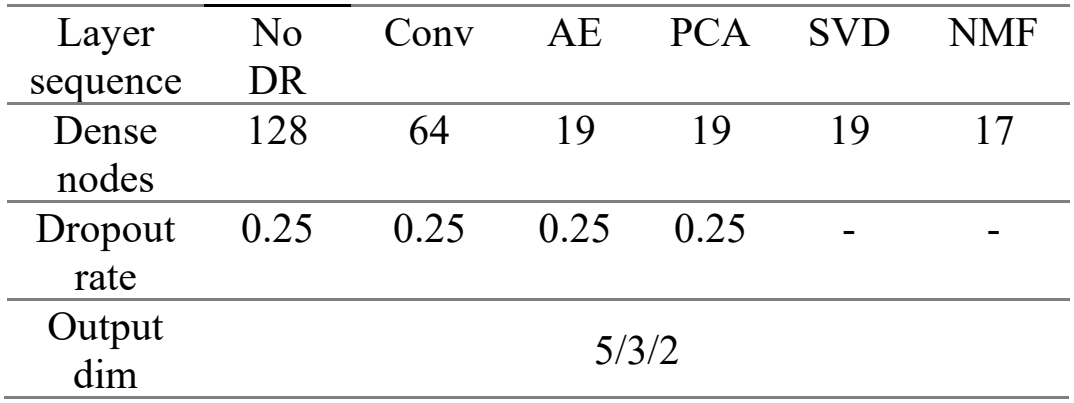
SNN ARCHITECTURE.

| Layer sequence | No DR | Conv | AE | PCA | SVD | NMF |
|---|---|---|---|---|---|---|
| Dense nodes | 128 | 64 | 19 | 19 | 19 | 17 |
| Dropout rate | 0.25 | 0.25 | 0.25 | 0.25 | - | - |
| Output dim | 5/3/2 |  |  |  |  |  |

**TABLE V.**
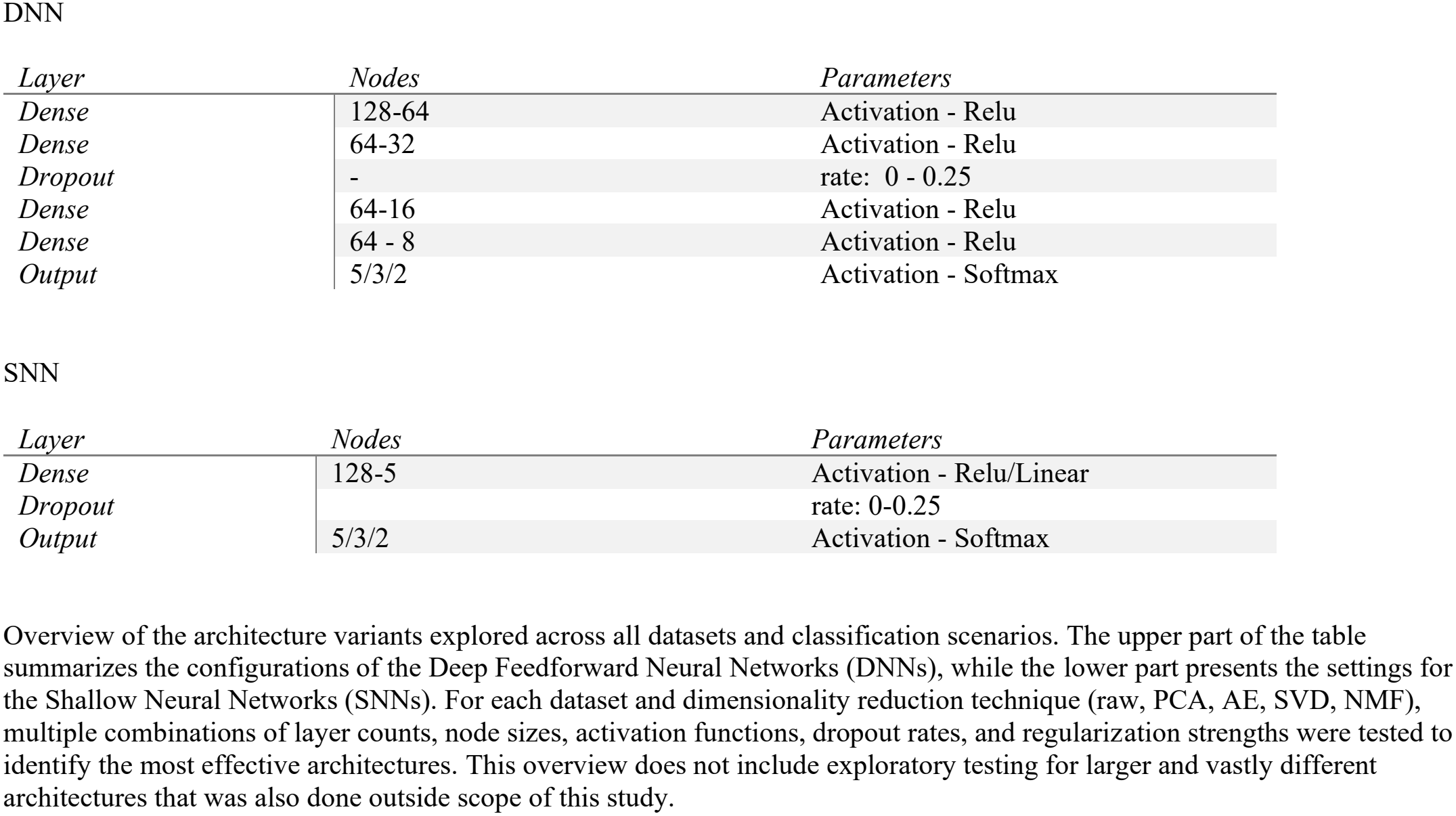
Tested Architecture parameters.

| DNN |  |  |
|---|---|---|
| *Layer* | *Nodes* | *Parameters* |
| *Dense* | 128-64 | Activation - Relu |
| *Dense* | 64-32 | Activation - Relu |
| *Dropout* | - | rate: 0 - 0.25 |
| *Dense* | 64-16 | Activation - Relu |
| *Dense* | 64 - 8 | Activation - Relu |
| *Output* | 5/3/2 | Activation - Softmax |
| SNN |  |  |
| *Layer* | *Nodes* | *Parameters* |
| *Dense* | 128-5 | Activation - Relu/Linear |
| *Dropout* |  | rate: 0-0.25 |
| *Output* | 5/3/2 | Activation - Softmax |

**TABLE VI.**
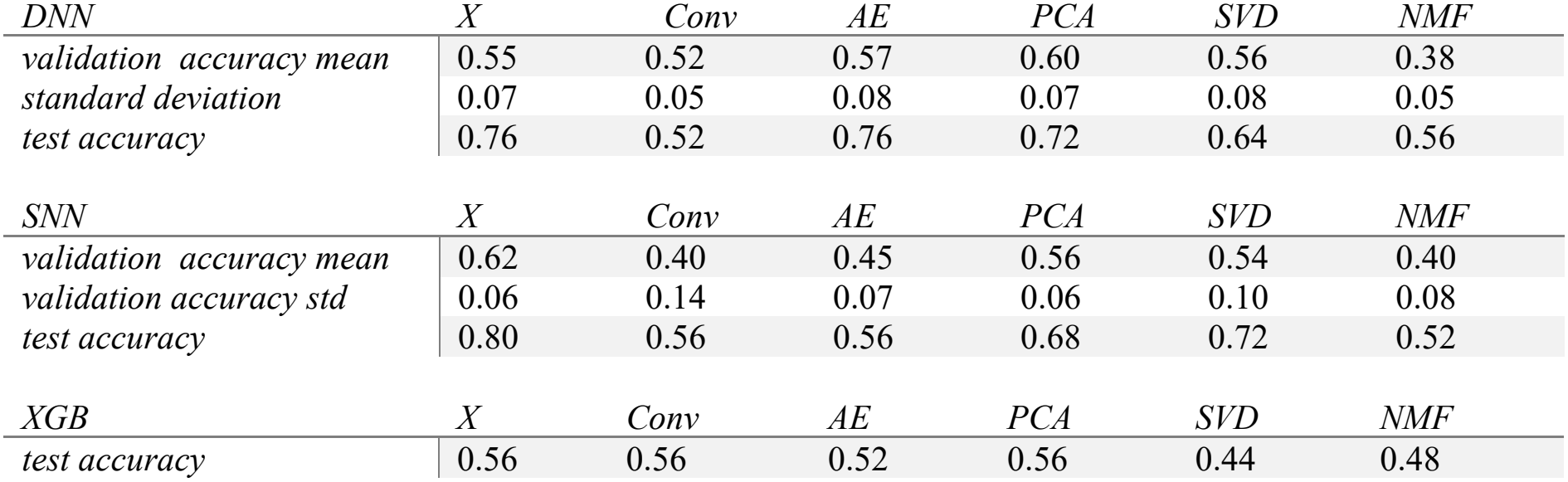
5-Class-Split Results.

| *DNN* | *X* | *Conv* | *AE* | *PCA* | *SVD* | *NMF* |
|---|---|---|---|---|---|---|
| *validation accuracy mean* | 0.55 | 0.52 | 0.57 | 0.60 | 0.56 | 0.38 |
| *standard deviation* | 0.07 | 0.05 | 0.08 | 0.07 | 0.08 | 0.05 |
| *test accuracy* | 0.76 | 0.52 | 0.76 | 0.72 | 0.64 | 0.56 |
| *SNN* | *X* | *Conv* | *AE* | *PCA* | *SVD* | *NMF* |
| *validation accuracy mean* | 0.62 | 0.40 | 0.45 | 0.56 | 0.54 | 0.40 |
| *validation accuracy std* | 0.06 | 0.14 | 0.07 | 0.06 | 0.10 | 0.08 |
| *test accuracy* | 0.80 | 0.56 | 0.56 | 0.68 | 0.72 | 0.52 |
| *XGB* | *X* | *Conv* | *AE* | *PCA* | *SVD* | *NMF* |
| *test accuracy* | 0.56 | 0.56 | 0.52 | 0.56 | 0.44 | 0.48 |

**TABLE VII.** 3-Class-Split Results.

| *DNN* | *X* | *Conv* | *AE* | *PCA* | *SVD* | *NMF* |
|---|---|---|---|---|---|---|
| *validation accuracy mean* | 0.70 | 0.72 | 0.67 | 0.69 | 0.70 | 0.56 |
| *validation accuracy std* | 0.10 | 0.02 | 0.07 | 0.09 | 0.08 | 0.07 |
| *test accuracy* | 0.84 | 0.52 | 0.84 | 0.80 | 0.80 | 0.84 |
| *SNN* | *X* | *Conv* | *AE* | *PCA* | *SVD* | *NMF* |
| *validation accuracy mean* | 0.70 | 0.60 | 0.67 | 0.68 | 0.62 | 0.61 |
| *validation accuracy std* | 0.06 | 0.10 | 0.05 | 0.07 | 0.05 | 0.14 |
| *test accuracy* | 0.88 | 0.60 | 0.76 | 0.80 | 0.60 | 0.80 |
| *XGB* | *X* | *Conv* | *AE* | *PCA* | *SVD* | *NMF* |
| *test accuracy* | 0.68 | 0.64 | 0.72 | 0.68 | 0.64 | 0.80 |

**TABLE VIII.** 2-Class-Split Results.

| *DNN* | *X* | *Conv* | *AE* | *PCA* | *SVD* | *NMF* |
|---|---|---|---|---|---|---|
| *validation accuracy mean* | 0.77 | 0.81 | 0.76 | 0.75 | 0.77 | 0.67 |
| *validation accuracy std* | 0.05 | 0.06 | 0.07 | 0.07 | 0.06 | 0.05 |
| *test accuracy* | 0.80 | 0.88 | 0.84 | 0.80 | 0.84 | 0.92 |
| *SNN* | *X* | *Conv* | *AE* | *PCA* | *SVD* | *NMF* |
| *validation accuracy mean* | 0.76 | 0.68 | 0.77 | 0.77 | 0.74 | 0.68 |
| *validation accuracy std* | 0.06 | 0.06 | 0.08 | 0.05 | 0.05 | 0.03 |
| *test accuracy* | 0.84 | 0.76 | 0.88 | 0.76 | 0.76 | 0.88 |
| *XGB* | *X* | *Conv* | *AE* | *PCA* | *SVD* | *NMF* |
| *test accuracy* | 0.84 | 0.84 | 0.80 | 0.72 | 0.84 | 0.80 |

**TABLE IX.** Software Used.

| **Package Name** | **Citation** | **Purpose of Use** |
|---|---|---|
| Keras (v3.3.3) | Chollet, F., & others. (2015). *Keras*. GitHub. Retrieved from https://github.com/fchollet/keras | Used as the high-level API on top of TensorFlow for creating, training, and testing deep learning models. |
| NumPy (v1.26.4) | Harris, C. R., Millman, K. J., van der Walt, S. J., Gommers, R., Virtanen, P., Cournapeau, D., … Oliphant, T. E. (2020). *Array programming with NumPy*. *Nature*, 585, 357–362. https://doi.org/10.1038/s41586-020-2649-2 | Used for data manipulation tasks such as preprocessing and input/output transformations. |
| Pandas (v2.2.2) | McKinney, W., & others. (2010). *Data structures for statistical computing in Python*. In *Proceedings of the 9th Python in Science Conference* (Vol. 445, pp. 51–56). | Used for data manipulation and transformation tasks. |
| Python (v3.12.10) | Van Rossum, G., & Drake, F. L. (2009). *Python 3 Reference Manual*. Scotts Valley, CA: CreateSpace. | Used as the primary programming language for all analyses in the study. |
| Scikit-learn (v1.5.0) | Pedregosa, F., Varoquaux, G., Gramfort, A., Michel, V., Thirion, B., Grisel, O., … others. (2011). *Scikit-learn: Machine learning in Python*. *Journal of Machine Learning Research*, 12(Oct), 2825–2830. | Used for dataset splitting, cross-validation, normalization, and dimensionality reduction via PCA. |
| Tensor Flow (v2.16.1) | Abadi, M., Barham, P., Chen, J., Chen, Z., Davis, A., Dean, J., … others. (2016). Tensorflow: A system for large-scale machine learning. In 12th USENIX Symposium on Operating Systems Design and Implementation (OSDI 16) (pp. 265–283). | Used as the backend framework for deep learning model |
| Matplotlib (v3.9.0) | J. D. Hunter, "Matplotlib: A 2D Graphics Environment," in Computing in Science & Engineering, vol. 9, no. 3, pp. 90-95, May-June 2007, doi: 10.1109/MCSE.2007.55. | For visualization |
| Seaborn (v0.13.2) | Waskom, M. L., (2021). seaborn: statistical data visualization. Journal of Open Source Software, 6(60), 3021, https://doi.org/10.21105/joss.03021. | For visualzation |
| XGBoost (v2.1.2) | T. Chen and C. Guestrin, "XGBoost: A Scalable Tree Boosting System," in Proceedings of the 22nd ACM SIGKDD International Conference on Knowledge Discovery and Data Mining, San Francisco, CA, USA, 2016, pp. 785–794. | For XGBoost baseline implementation |

The same as for embedding, also for the classification networks, hyperparameter tuning was also conducted empirically. While not all tested configurations are reported in the main text, a wide range of architectural variants consisting of variations in layer compositions, node counts and various activation functions as well as other hyperparameters, was explored (Appendix, Table V).

### D. Simulation Study

To systematically evaluate the statistical properties of the proposed classification approaches, a simulation study was conducted. The simulated data sets were based on the CLR-transformed original dataset and retained its dimensions (125 × 125) consisting of five classes (25 samples each). In the first step, an empirical standard deviation vector 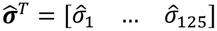, with 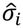 representing an estimated standard deviation of feature *i* from the control group was computed and randomly permuted. The simulated control dataset was then generated by drawing 125 features for 25 subjects, with normalized counts for each subject sampled from a normal distribution 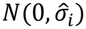. Experimental groups were generated using a mean-shift parameter (*S*). Each simulated experimental set consisted of *P*0 features with normalized counts sampled from 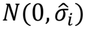 and *Ps* = (125 − *P*0) features represented bacterial families influenced by supplementation with normalized counts sampled as follows:

- 25 individuals representing class 1: 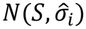,
- 25 individuals representing class 2: 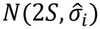,
- 25 individuals representing class 3: 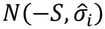,
- 25 individuals representing class 4: 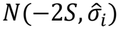.

The number of affected features (*Ps*) ranged from 5 to 125 with a step size of 5, the mean-shift parameter (*S*)ranged from 0.1 to 2.0 with a step size of 0.1. Each simulation scenario was repeated ten times for every parameter combination. For classification, the models architecture and DRT approaches were kept the same as for case study. Model classification accuracy was averaged across the ten simulations. The relationship between test accuracy, *Ps*, and *S* was assessed as Spearman correlation coefficient between accuracy and one of the simulation parameters (*S*), as well as by fitting the polynomial regression model for visual comparison.

## IV. Results

### A. Data Inspection

The initial step of data pre-processing revealed extreme zero-inflation (sparsity) in the bacterial family abundance matrix. In particular, 64 bacterial families (50%) were detected in only 6 samples (<5% of the total), whereas only 4 families (3%) were detected in at least 95% of samples (118 samples) (Fig. 4). Moreover, the distribution of read counts was highly imbalanced across individual fish and bacterial families (Fig. 5). The majority (67%) of features (85 families) were represented by less than 100 reads, while a small subset of features showed extremely high abundance, with the most extreme case of 3 families (2%) represented by over 10,000 reads each.

**Fig. 4.**
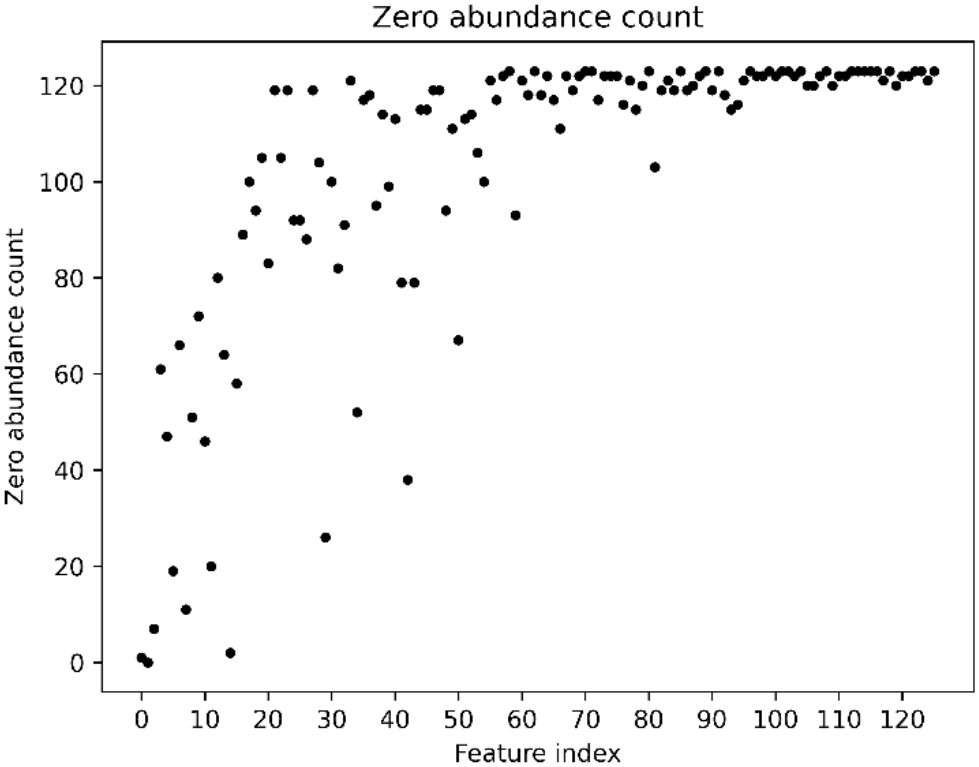
Zero abundance distribution. Shows many features (50%) were present in small number (5%) of samples only

**Fig. 5.**
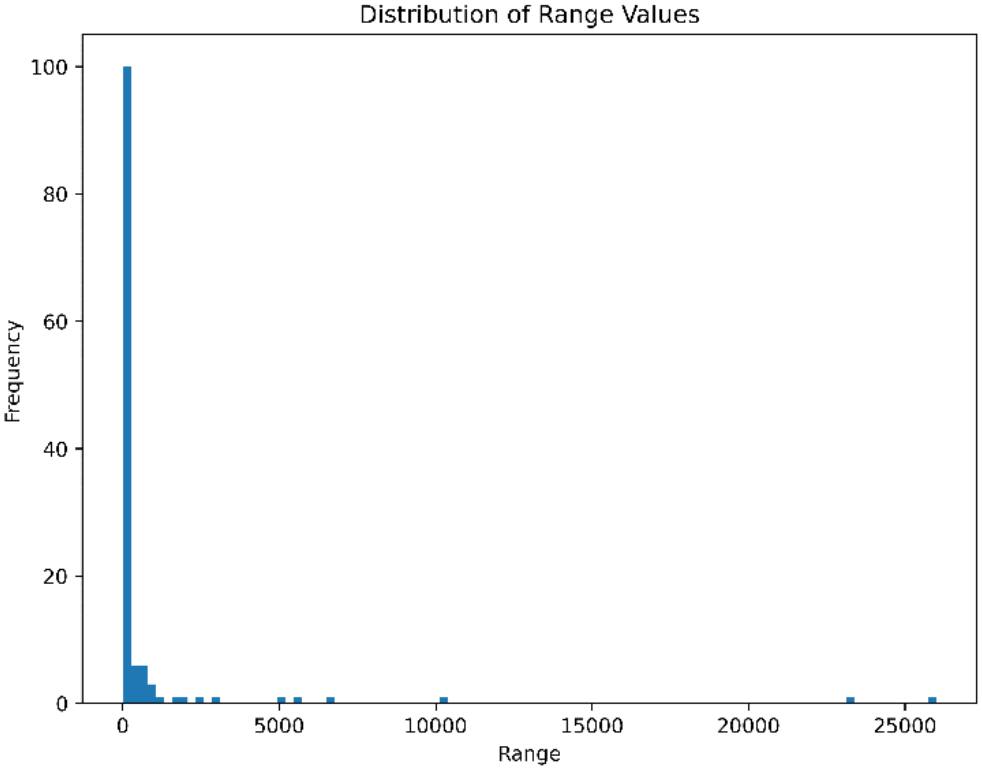
Distribution of raw read counts. Many features had similar range of counts(0-100) while some significantly differed (>10000)

### B. Case Study

#### Five-class split

The objective was to discriminate between all experimental conditions (Control and experimental Groups 1–4) for which the expected by chance classification accuracy is 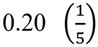. For DNN, cross-validation (CV) accuracies remained relatively consistent across most dimensionality reduction techniques, ranging from 0.52 ± 0.05 for the Conv approach to 0.60 ± 0.07 for PCA, with the most underperforming embedding being NMF (0.38 ± 0.05). Test accuracies exhibited a higher variation. The best performance (0.76) was achieved by both AE and no dimensionality reduction, whereas the lowest accuracies were recorded for Conv (0.52) and NMF (0.56). For SNN, cross-validation accuracies demonstrated high variability. The best accuracy was achieved for the no dimensionality reduction approach (0.62 ± 0.06), whereas the worst classifiers used Conv (0.40 ± 0.16) and NMF (0.40 ± 0.08) embeddings. The highest test accuracy of SNN was for the unreduced feature space (0.80), with NMF consistently yielding the lowest accuracy of 0.52. The XGBoost benchmark demonstrated consistent classification quality, achieving a best test accuracy of 0.56 for no reduction, Conv, and PCS scenarios, and the lowest accuracy of 0.44 for SVD.

Standard deviation (SD) of CV differed substantially across both architectures and DRT combinations, ranging from 0.05 to 0.14 (Fig. 6, Appendix Table VI). SNN, especially for Conv and NFM embeddings, showed consistently higher SDs of accuracy across CV evaluations, indicating low model robustness.

**Fig. 6.**
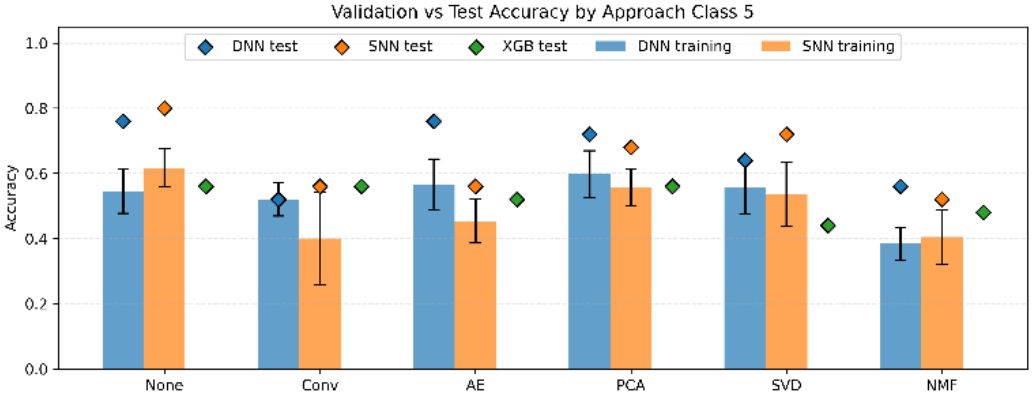
Classification performance across dimensionality reduction techniques in the five-class setting. Evaluated architectures are color-coded: blue for Deep Neural Network (DNN), orange for Shallow Neural Network (SNN), and green for XGBoost. Bars represent mean cross-validation accuracy (whiskers: standard deviation); points represent test set accuracy. The random assignment baseline is 0.20. Test set accuracies generally exceeded cross-validation means and XGBoost baselines across models. The highest test accuracy was achieved by unreduced SNN (0.80), followed by unreduced DNN and AE-DNN (0.76), while NMF-SNN and Conv-DNN yielded the lowest test scores (0.52).

#### Three-class split

A three-class classification scenario discriminated the following conditions: Class 0 (Control), Class 1 (combined Groups 1 and 3 – only water supplementation), and Class 2 (combined Groups 2 and 4 – feed and water supplementation). The expected by chance classification accuracy was 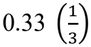 in this scenario. For DNN, mean CV accuracies remained stable across most dimensionality reduction techniques, ranging from 0.72 ± 0.02 for Conv embedding to 0.56 ± 0.07 for NMF. Test set accuracies were overall higher, reaching 0.84 for no DRT, AE, and NMF, followed by PCA and SVD (0.80) and 0.52 for Conv. For SNN, mean validation accuracies ranged from 0.60 ± 0.10 for the convolution-based approach to 0.70 ± 0.06 for no reduction. Test results exhibited higher dispersion: unreduced SNN achieved the highest accuracy of 0.88, while convolution-based embedding and SVD resulted in the lowest performance (0.60). The XGBoost benchmark achieved test accuracies between 0.64 (Conv, SVD) and 0.80 (NMF), representing the best classification performance across all scenarios.

For the three class split, the SNN architecture was generally more stable than DNN with SD ranging from 0.5 to 0.7, except for Conv and NMF embeddings (0.10 and 0.14, respectively). Whereas for 5-class split DNN’s usually exhibited lower CV-SD (Fig 7, App Table VII).

**Fig. 7.**
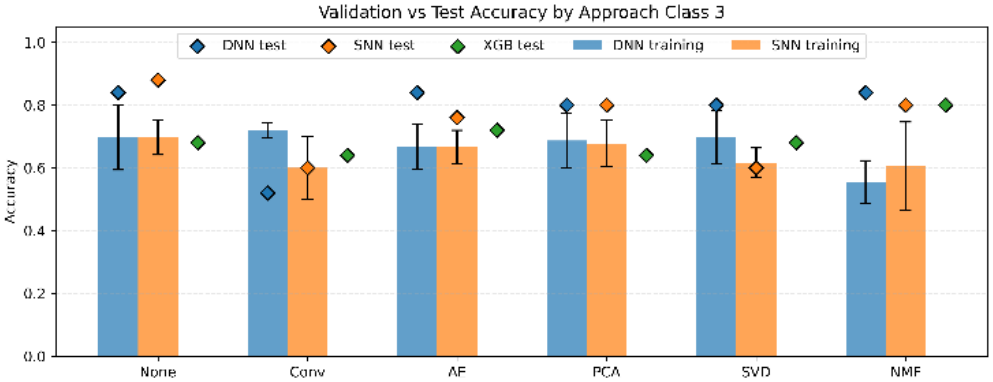
Classification performance in the three-class setting across dimensionality reduction techniques. Overall classification performance improved relative to the five-class task, with nearly all model configurations surpassing the 0.33 random baseline. The unreduced SNN model achieved the highest overall test accuracy (0.88), followed closely by the unreduced, AE, and NMF DNN models (all 0.84). Notably, NMF feature extraction substantially boosted XGBoost performance to its overall peak of 0.80, 1D-convolution-based architectures consistently yielded the lowest test accuracies.

#### Two-class split

A binary classification scenario evaluated discrimination between lack (combined control, Groups 1 and 3) and presence of feed supplementation (combined Groups 2 and 4). In this setting, the random chance baseline accuracy is 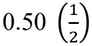. For DNN, cross-validation accuracies ranged from 0.67 ± 0.05 for NMF to 0.81 ± 0.06 for Conv. Test accuracy was high across all configurations, with the lowest accuracies of 0.80 estimated for the no reduction scenario and PCA, while the highest accuracy of 0.92 was obtained for NMF - the highest accuracy across all setups in our study. For SNN, cross-validation accuracies demonstrated high consistency, ranging from 0.68 ± 0.06 for Conv to 0.77 ± 0.08 for AE and 0.77 ± 0.05 for PCA. Test accuracies ranged from 0.76 for Conv, PCA, and SVD embeddings to 0.88 for NMF and AE. The XGBoost benchmark yielded 0.84 classification accuracies across the no reduction scenario, Conv, and SVD, dropping to 0.72 for PCA. The standard deviation of cross-validation runs remained consistently low across scenarios (0.05 to 0.08), except for SNN with NMF embedding, which exhibited the highest consistency (0.03), indicating the best loss optimization behavior in the binary classification task (Fig 8, App. Table VIII).

**Fig. 8.**
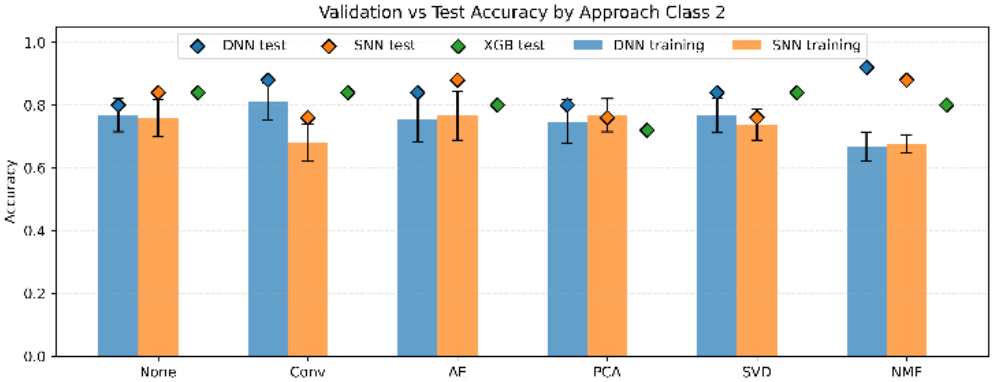
Classification performance in the binary (two-class) setting across dimensionality reduction techniques. Binary classification yielded the strongest overall predictive performance, with all model configurations substantially exceeding the 0.50 random baseline. The NMF-DeepNN model achieved the highest test accuracy (0.92) recorded across the entire study, while NMF and AE features enabled SNN models to reach 0.88 test accuracy. Unlike the multi-class scenarios where 1D-convolution features underperformed, the convolution-based DNN model demonstrated high validation accuracy (0.81 ± 0.06), and NMF-SNN displayed the highest overall training stability.

### C. Simulation Study

The simulation study demonstrated a significant (*p* < 0.05) positive correlation (0.59) between the mean shift parameter (*S*) and test accuracy. As effect size increased, accuracy increased from the range of 0.20 - 0.40 at *S* = 0.1 to 0.40 - 1.00 at *S* = 0.6 (Fig. 9A), beyond which accuracy leveled off. An exception to this trend occurred for the lowest number of affected features (*Ps* = 5), where accuracy remained in the range of 0.20 - 0.80, suggesting that very sparse signal is hinders classification performance, regardless of effect size. Overall, the number of affected features exhibited only a weak and non-significant relationship with test accuracy (0.34). For most values of *Ps*, accuracy remained stable with a single exception of the aforementioned drop at *Ps* = 5 (Fig. 9B).

**Fig. 9.**
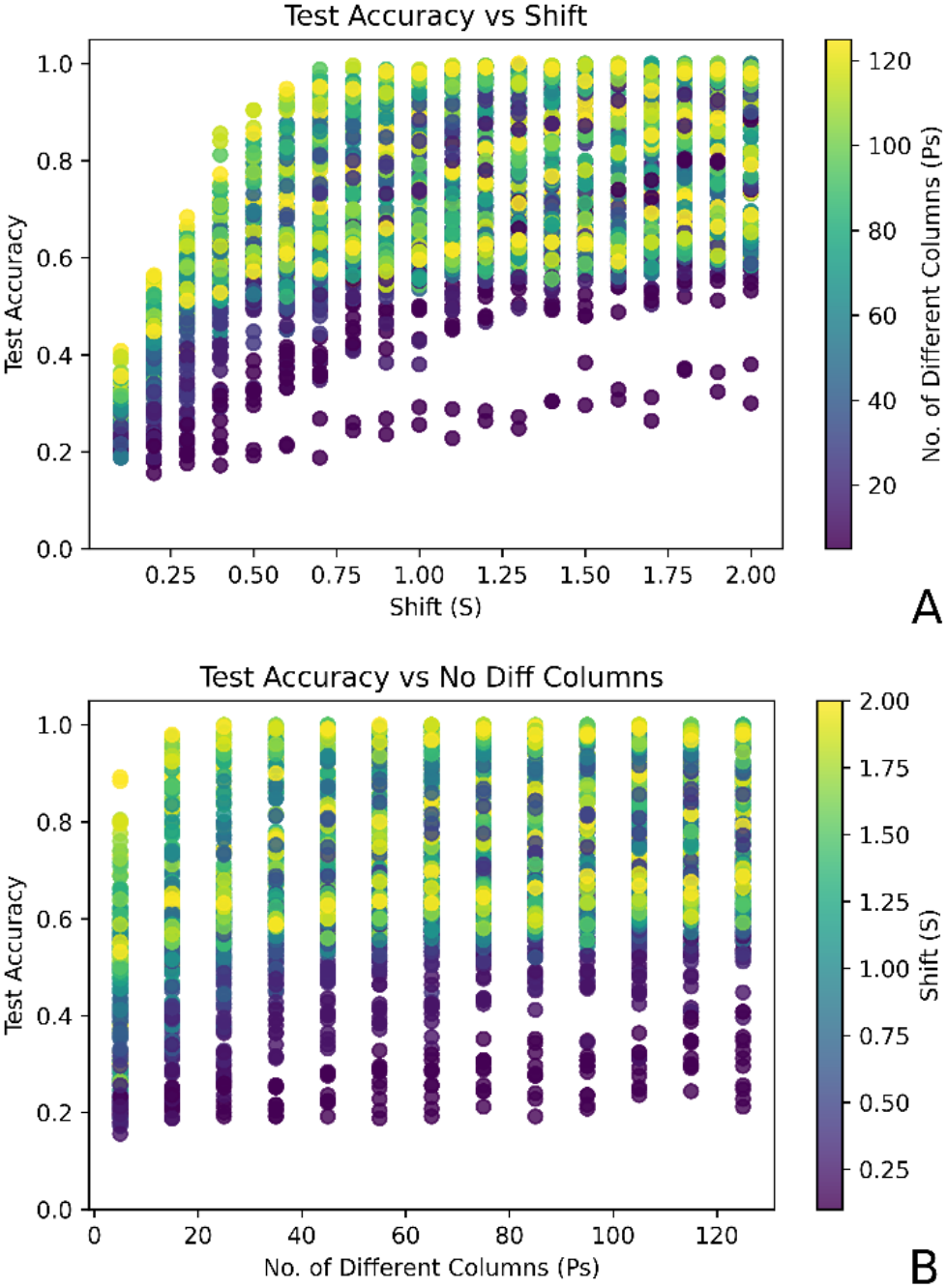
Simulation study results comparing test accuracy against the mean shift parameter (*S*) and the number of affected features (*Ps*). (A, top) A strong positive relationship is observed between test accuracy and *S*. (B, bottom) In contrast, test accuracy shows minimal dependence on *Ps*, except at the lowest threshold (*Ps* = 5), which exhibits a notable decrease (~0.20) in maximum accuracy.

Fitting a second-order polynomial regression of test accuracy on the effect size (*R*^2^ = 0.6) demonstrated that DNN architectures consistently outperformed SNN across all mean-shift scenarios (Fig 10). The highest performance was observed for PCA and SVD embeddings, which reached an accuracy of 0.73 already at *S* = 0.5. The next best results at this mean-shift (0.63) were from no DRT and AE embedding, while the Conv reduction approach performed the worst (0.53) among the DNN models. These differences remained consistent at higher *S*. Note that, unlike other scenarios, the observed accuracy of AE embedding did not leveled-off but monotonically increased with *S*. SNN architectures demonstrated lower overall performance.

**Fig 10.**
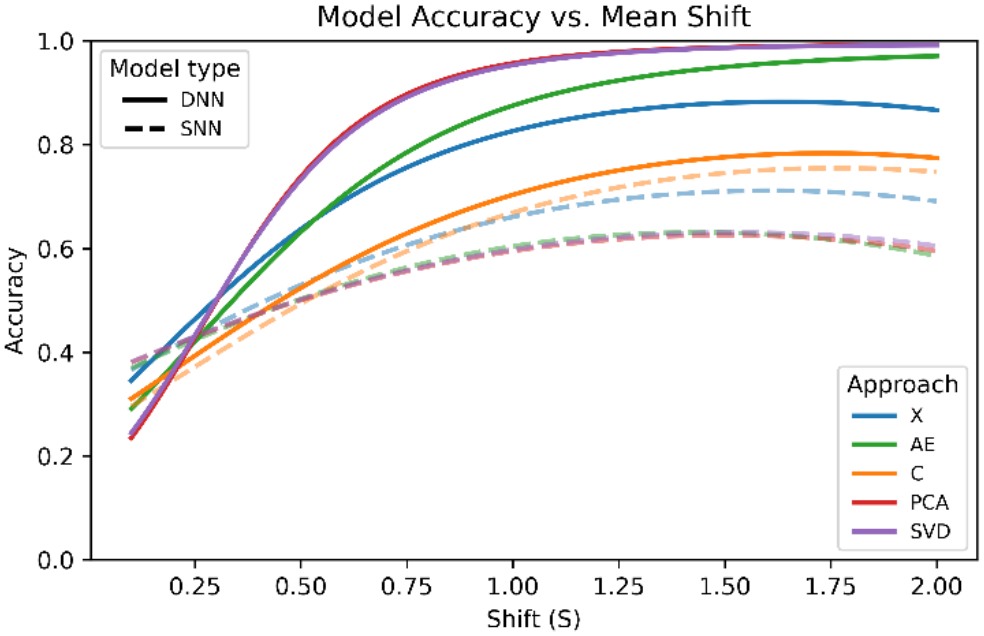
Polynomial regression models fitted to various classification approaches. Highlighted Approaches represent Deep Neural Networks, While the perforated represent Shallow Neural Network.

The simulation study revealed a significant, moderately positive correlation (0.22) between computational time and the mean shift, with an even stronger correlation (0.42) between computational time and the number of affected features. Computational time was primarily dependent on DRT. AE embedding was the most computationally intensive, with a mean runtime of approximately 113 seconds per simulation. This is nearly four times higher than the no reduction baseline, which averaged only 29 seconds per simulation. Also the number of training epochs required for convergence exhibited a positive correlation with computational time (0.38), which was heavily influenced by network depth. SNNs required fewer epochs, ranging from 70 for the convolution-based embedding to 100 for other DRTs. DNNs required substantially more epochs, ranging from 192 for the no reduction scenario to 270 epochs for PCA and SVD. Notably, the DNN architecture with Conv converged in only 66 epochs.

## V Conclusions

### A. Case Study

The observed improvement in classification accuracy as the number of classes decreases can be attributed to both methodological and biological factors. Methodologically, reducing the number of classes simplifies the decision boundary and thus diminishes the potential of misclassification. Biologically, grouping classes may better reflect the underlying biological process (here the microbial community dynamics).

Across our evaluations, no single architecture (DNN or SNN) demonstrated superiority consistent across all dimensionality reduction techniques and class splits. However, on the unreduced feature space, despite lower model capacities expressed by the number of weighs, SNN performed at least as well as DNN. This suggests that raw data representations may lack the structural complexity required to leverage the higher informational capacity of deeper architectures (i.e does not contain enough information to estimate a larger number of weights) and thus increasing susceptibility to fitting fake patterns to noise, within a small sample size. Generally, models trained on unreduced data achieved comparable or superior performance relative to those using dimensionality reduction. This finding indicates that dimensionality reduction is not universally beneficial and in small data sets may even lead to reduced classification. The exception were convolution-based feature representations, for which DNN consistently outperformed shallow models, suggesting that convolutional transformations results in feature spaces that are better suited for higher-capacity models. However, the overall peak accuracy of 0.92 was achieved using non-negative matrix factorization (NMF) combined with a deep neural network in the binary setting. Neural network classifiers generally outperformed the XGBoost baseline, demonstrating that even with restricted sample sizes, neural networks are more flexible methods to effectively capture patterns.

Models trained on unreduced data exhibited the highest robustness across classification scenarios, maintaining stable performance across all three classification scenarios. That is not necessarily an advantage, as the underlying robustness hindered the decision of which pattern best reflects the underlying biological situation. In contrast, NMF demonstrated high sensitivity to class resolution, performing exceptionally well in simpler classification tasks but suffering noticeable performance degradation in the 5-class scenario. This sensitivity of NMF offers insights into the true biological structure of the data.

Additionally, the observation that accuracies from models trained on validation and tested on test set, frequently exceeded cross-validation mean accuracies is likely attributable to the larger effective sample size available during final model fitting on the combined training and validation sets. Finally, while the dataset severe sparsity and zero-inflation increased the risk of overfitting, incorporating dropout layers and *L*_2_ regularization proved very helpful for overfitting prevention and thus increasing model training stability. Nonetheless, persistent variability across experimental runs underscores the challenge of model robustness in small-sample biological data.

### B. Simulation Study

The number of affected features *Ps* had a negligible impact on classification performance, unless *Ps* was very low, indicating that only a few features differ in abundance between groups. With *Ps* = 5, classification accuracy was substantially degraded, demonstrating that even under large class separations expressed by mean shift (*S*), an insufficient number of discriminatory features fails to provide the support for classification. This is furthermore supported by the result that *S* exerted a notably stronger effect on accuracy, confirming that effect size serves as the primary determinant of classification quality. Under simulated conditions, Deep Neural Networks consistently outperformed shallow models achieving high accuracy exceeding 0.95. This demonstrates DNN superior capacity to capture complex patterns when the underlying signal is sufficiently strong.

PCA and SVD were the top approaches among the evaluated dimensionality reduction techniques, autoencoders yielded moderate results, and convolution-based approaches resulted in the lowest accuracies. This similar performance of PCA and SVD stems from their algebraic equivalence on mean-centered, normally distributed data, where both methods compute orthogonal projections that maximize global variance capture. This indicates that preserving global linear dependencies is more advantageous in this context than extracting local, non-linear patterns However, the relative underperformance of the AE and Conv embeddings highlights a trade-off between a parameter-free, deterministic PCA and SVD and deep representation models sensitive to hyperparameters - such as latent space size in AE or kernel size in Conv, which was not formally explored in the current study.

Computational cost increased more with the number of affected features (*Ps*) than with the effect size (*S*). Autoencoders proved to be the most computationally expensive due to the requirement of training a separate DL architecture just for feature extraction. Training time increased with the number of epochs and varied considerably across model types. DNN architectures required substantially more training epochs and computational time than SNN, a consequence of their larger number of trainable parameters. Certain embeddings, particularly Conv, demonstrated faster convergence but suffered from lower classification accuracy.

### C. Outcome

Taken together, the case study and simulation results provide a cohesive explanation for model performance in microbiome classification tasks. The simulation demonstrated that classification accuracy is primarily driven by signal strength rather than the number of affected features, and that Deep Neural Networks perform well at extracting complex patterns when sufficient signal exists. This insight contextualizes the case study results, where performance improved alongside reduced class complexity, a shift that likely reflects both increased effective signal strength and simplified decision boundaries. Additionally, the simulation findings indicated that PCA and SVD effectively preserve informative global structures, aligning with their strong, consistent performance on real data. In contrast, convolution-based approaches underperformed during simulations, implying that structural pattern extraction is less suited for this data type, a trend partially observed in the case study. Nonetheless, differences between simulated and empirical data, (i.e. sparsity and distribution) likely account for the observed discrepancies between the two settings.

A critical insight from the case study is that models trained on unreduced data exhibited strong robustness across varying class splits, whereas certain dimensionality reduction techniques, such as NMF, displayed pronounced sensitivity to task complexity. Looking from the perspective of the simulation results, this suggests that specific transformations may artificially amplify or suppress signals depending on class structure, rendering their effectiveness highly context-dependent rather than universal. Although the simulations established a clear superiority for deep neural networks, this advantage was only partially observed in the case study, where deep models showed no consistent edge over their shallow counterparts. This discrepancy likely stems from the limited sample size, high sparsity, and weak signal inherent to the real dataset, all of which constrain the ability of deep models to fully utilize their representational capacity while simultaneously increasing the risk of overfitting. Ultimately, both studies show a fundamental trade-off. While Deep learning models offer exceptional pattern extraction capabilities under favorable conditions, they demand substantially larger computational resources and larger sample sizes which is not usually the case in biological or medical studies. In the context of small, sparse microbiome datasets, predictive performance remains sensitive to data representation and signal characteristics, highlighting the importance of preprocessing and method selection.

## VI. Discussion

This study focused on a methodological exploration of the classification performance of various dimensionality reduction techniques coupled with simple dense neural network-based classifiers for small count data, with class unambiguity and the number of features similar to the number of samples representing a mild version of P>>N problem.

### A. Data inspection and preprocessing

Initial data exploration revealed a high proportion of zero abundance cells. From a statistical perspective, such zero-inflation not only violates the parametric distributional assumptions underlying classical inference methodologies, but also presents a computational problem when zeroes are involved in calculations. Biologically, such sparse structure is characteristic of many data types, such as microbiome or protein abundance, gene or transcript expression, especially in single cell transcriptomics, or many covariance matrices. In microbiome data, which was used here as the biological example, zero-abundance either represents taxa truly absent in the analyzed environment or taxa that are abundant but remain below the detection sensitivity of the sequencing method. Despite the challenges, several machine learning architectures evaluated in this study achieved high classification quality, indicating that zero inflation did not significantly compromise predictive power, which is consistent with the findings of Haghani et al. [31].

From the perspective of DL, the extreme variance in feature abundance across samples introduces the risk of numerical instability in computations, as well as the problem of highly abundant taxa disproportionately dominating the learning process. To address the problem of zero-inflation, the CLR transformation was applied at data pre-processing step, reducing variance range differences and helping to ensure that all bacterial families, regardless of abundance, contribute equitably to the learned latent representations [32]. Moreover, the CLR transformation has been shown to improve the performance of not only DL models but also classical parametric statistical methods [5], [33].

### B. Dimensionality reduction and model performance

In the preliminary stages of this study, other (nonlinear) dimensionality reduction techniques, such as t-SNE and UMAP, were also considered [34]. However, they were excluded due to their poorer convergence, and sensitivity to hyperparameter tuning. Given the need to simultaneously evaluate both dimensionality reduction techniques and DL architectures, it was important to use a DRT approach that estimates deterministic and reproducible embeddings. This allowed the focus to remain on assessing the performance of the neural network models without introducing additional variability from, less consistent dimensionality reduction methods [35], [36]. Furthermore, although applying DRTs generally accelerates downstream neural network convergence – as demonstrated by Velliangiri et. al. [37], our results indicated that the computational overhead of calculating embeddings often negated possible optimization benefits. This, however, might be attributable to the small overall dimensions of the simulated and demonstration data sets. Moreover, the benefit of DRT may depend on the inter-feature correlation structure. In the case study data set correlation between features varied with some highly correlated (−0.64, +0.99, Pearson) while the others showed insignificant correlation. As for the simulation study, features were generated as uncorrelated.

The maximum 0.92 accuracy was achieved for a test dataset. This suggests the presence of outlier samples or feature counts, and it is tempting to state that improving test accuracy beyond this level would require models with greater capacity, represented by deeper and more highly parameterized architectures, to capture more complex relationships within the data, our observation was that such model would require much larger dataset which was not the case in this study and often is not feasible in biological studies. Additionally, attempting to force higher accuracy by fitting very deep, highly parameterized architectures, would likely exacerbate overfitting rather than improve generalization. Another important aspect that would help to alleviate accuracy would be to use auxiliary features or to formally model phylogenetic similarity between microorganisms. One particularly interesting approach to the latter was proposed by Nguyen et. al. [38], where phylogenetic trees were constructed and visualized as images with pixel density representing bacterial abundance in each taxon. These images were then classified using convolutional neural networks. While promising, such an approach would require substantially larger datasets to avoid overfitting, as well as precise taxonomic annotation of features, which for S16 rRNA sequencing data is limited to a family or, in the best case, to the genus level. In our study, overfitting was frequently observed in CNN-based configurations, likely due to their high representational capacity. In small-sample contexts, highly parametrized models tend to learn noise in addition or instead of a true signal. This was confirmed by our observation on CNN performance with sparse signal for simulated data (*Ps* = 5) where it’s underperformance was more profound than other DRT’s suggesting it’s susceptibility for noise fitting. This underscores the critical need to match model complexity to dataset dimensions. In this context, dimensionality reduction techniques serve as a valuable tool for compressing the input features into a lower-dimensional manifold that is supposed to retain only the essential discriminatory information. By reducing input dimensionality while retaining essential information, DRTs may help more complex models to generalize effectively, even when the data is small [39].

### C. Computational Efficiency and Scalability

All our experimental configurations converged within 15 minutes per pipeline on a standard consumer-grade PC, indicating that computational resource demands are minimal for microbiome datasets of this scale. Peak RAM allocation did not exceed 2.5 GB. Among the dimensionality reduction techniques, a deterministic PCA was considerably faster to compute than AE based embeddings, as it involved training the whole AE. Still, computational scalability remains an important consideration for larger datasets.

### D. Limitations

A primary limitation of this study is the omission of feature importance analysis, which limits the biological interpretability of the models. Since the microbiome dataset was used as a demonstrative biological set of count data, we deliberately restricted ourselves to methodological aspects rather than the extraction of biological information that has already been explored elsewhere [40]. Methodologically, we imposed a constraint to evaluate only dense neural network based classifiers. Consequently, we did not explore the whole space of alternative deep learning models, for example, approaches proposed by Chen et. al. [41] and Maryam et.al. [42] based on graph and recurrent neural networks or transformer architectures [43]. This was related to the major objective of our study to explore the potential of very simple and shallow configurations to discriminate samples between experimental conditions under the small sample, P>>N, and unknown number of classes problem.

As for dimensionality reduction we are well aware that from the plethora of possible dimensionality reduction approaches and their architectures only a small subset was considered in this study. As it is not possible and even not meaningful to test multitude of dimensionality reduction and embedding methods, we compared approaches that we believe extract different aspects of underlying feature maps. Such as: deterministic PCA and SVD approaches that utilize the feature covariance and feature values matrices, respectively; deterministic NFM that is dedicated for sparse data; semi-stochastic DL approaches that estimate embeddings as latent space representations of features (AE) or through the convolution operation (1DConv).

## Data and code availability

The 16 S rRNA Gene Sequencing Data is available in the NCBI BioProject repository. It can be accessed using the following DOI or accession ID: https://identifiers.org/ncbi/bioproject:PRJNA1315210. Code and Feature Table data used in this study is available in repository under link: https://github.com/ThetaUP/Does-embedding-impact-classification-performance-A-small-sample-perspective

## Supplementary Materials

